# A modular protein subunit vaccine candidate produced in yeast confers protection against SARS-CoV-2 in non-human primates

**DOI:** 10.1101/2021.07.13.452251

**Authors:** Neil C. Dalvie, Lisa H. Tostanoski, Sergio A Rodriguez-Aponte, Kawaljit Kaur, Sakshi Bajoria, Ozan S. Kumru, Amanda J. Martinot, Abishek Chandrashekar, Katherine McMahan, Noe B. Mercado, Jingyou Yu, Aiquan Chang, Victoria M. Giffin, Felix Nampanya, Shivani Patel, Lesley Bowman, Christopher A. Naranjo, Dongsoo Yun, Zach Flinchbaugh, Laurent Pessaint, Renita Brown, Jason Velasco, Elyse Teow, Anthony Cook, Hanne Andersen, Mark G. Lewis, Danielle L. Camp, Judith Maxwell Silverman, Harry Kleanthous, Sangeeta B. Joshi, David B. Volkin, Sumi Biswas, J. Christopher Love, Dan H. Barouch

## Abstract

Vaccines against SARS-CoV-2 have been distributed at massive scale in developed countries, and have been effective at preventing COVID-19. Access to vaccines is limited, however, in low- and middle-income countries (LMICs) due to insufficient supply, high costs, and cold storage requirements. New vaccines that can be produced in existing manufacturing facilities in LMICs, can be manufactured at low cost, and use widely available, proven, safe adjuvants like alum, would improve global immunity against SARS-CoV-2. One such protein subunit vaccine is produced by the Serum Institute of India Pvt. Ltd. and is currently in clinical testing. Two protein components, the SARS-CoV-2 receptor binding domain (RBD) and hepatitis B surface antigen virus-like particles (VLPs), are each produced in yeast, which would enable a low-cost, high-volume manufacturing process. Here, we describe the design and preclinical testing of the RBD-VLP vaccine in cynomolgus macaques. We observed titers of neutralizing antibodies (>10^4^) above the range of protection for other licensed vaccines in non-human primates. Interestingly, addition of a second adjuvant (CpG1018) appeared to improve the cellular response while reducing the humoral response. We challenged animals with SARS-CoV-2, and observed a ~3.4 and ~2.9 log_10_ reduction in median viral loads in bronchoalveolar lavage and nasal mucosa, respectively, compared to sham controls. These results inform the design and formulation of current clinical COVID-19 vaccine candidates like the one described here, and future designs of RBD-based vaccines against variants of SARS-CoV-2 or other betacoronaviruses.

## Introduction

Prophylactic vaccination is effective in eliciting protective immunity against SARS-CoV-2 and preventing coronavirus disease 2019 (COVID-19) (*1*). Multiple vaccines have now been distributed at large scale in many countries, and have resulted in a lower incidence of infection and severe disease caused by SARS-CoV-2 (*2, 3*). Access to vaccines remains limited, however, in low- and middle-income countries (LMICs), where infectious variants of SARS-CoV-2 continue to emerge in large scale outbreaks (*4*). In addition to financial and logistical support from developed countries and health organizations, vaccines produced by local manufacturers could enable the lowest costs for interventions in these countries and potentially minimize the infrastructure required for their distribution (*5–7*). Protein subunit vaccines are a promising solution because they can be manufactured using existing large-scale microbial fermentation facilities in LMICs (*8*), typically do not require frozen storage and distribution, and are safe and effective when used with adjuvants (*9, 10*). Here, we describe the design and immunogenicity of a modular protein subunit vaccine, comprising a SARS-CoV-2 spike protein subunit receptor binding domain (RBD) displayed on a Hepatitis B virus-like particle (VLP) that is constructed using a covalent peptide-mediated linkage (SpyTag/SpyCatcher). Both of these vaccine components are currently produced by microbial fermentation at a large-scale manufacturing facility in India. We show that this vaccine candidate elicits a strong immune response in cynomolgus macaques and protects against SARS-CoV-2 challenge. Based on these promising data, this vaccine candidate is currently being tested in clinical trials (ANZCTR Registration number ACTRN12620000817943).

## Results

### Design of an accessible protein subunit vaccine

We sought to design a protein subunit vaccine that would be both suitably immunogenic and simple to manufacture for affordable distribution in LMICs. Multiple protein vaccines based on the trimeric SARS-CoV-2 spike protein have demonstrated efficacy, but are manufactured in insect or mammalian cells, which are difficult to transfer to existing facilities in LMICs (*11, 12*). The receptor binding domain (RBD) of the spike protein has been proposed as an alternative to the full spike protein because it has been shown to elicit multiple potent neutralizing antibodies directed at multiple epitopes (*13–15*), and can be manufactured in microbial systems like the biotechnological yeast *Komagataella phaffii* (*Pichia pastoris*) (*16, 17*). Formulations comprising only monomeric RBD (Wuhan-Hu-1) and adjuvant tested to date have required three doses to elicit potent neutralizing responses in humans (*18, 19*). While further optimization of such formulations could improve these designs, we and others have demonstrated that multimeric display of RBD on VLPs can be highly immunogenic (*20–24*).

Here, we selected Hepatitis B surface antigen (HBsAg) virus-like particles (VLPs) as a nanoparticle on which to display the RBD. This choice leverages extensive experience with a previously tested commercial product, GeneVac-B, that is manufactured at low cost and distributed in LMICs for prevention of Hepatitis B (*25*). As a model for our design here, we referenced previously reported designs with this core nanoparticle decorated with a malarial subunit antigen (*26*). A polypeptide-based system (SpyTag/SpyCatcher) allowed for covalent linkage of the antigen to the VLP by a transpeptidation reaction. This general design can increase antigen-specific antibody titers in mice, and the responses elicited in prior examples were unaffected by the presence of pre-existing antibodies against HBsAg (*27*). The modularity of the SpyTag/SpyCatcher system allows each component of the final particle to be expressed and purified independently to maximize yields and quality.

We adapted this approach to make a vaccine candidate for COVID-19. We genetically fused the SpyTag peptide onto the SARS-CoV-2 RBD. This fusion protein was manufactured in an engineered strain of *K. phaffii* (*28*). The RBD-SpyTag and HBsAg-SpyCatcher VLPs were each purified separately and then conjugated in a GMP process to produce the RBD-VLP antigen (Fig. 1A). In this study, the RBD-VLP antigen was formulated with two adjuvants: 1) aluminum hydroxide (alum) and 2) alum combined with CpG1018—a potent commercial TLR9 agonizing adjuvant known to elicit Th1-like responses (*29*). Analysis of the formulated vaccine drug product by SDS-PAGE showed only small fractions (<20%) of unconjugated HBsAg-VLP and RBD, and complete adsorption of the RBD-VLP antigen onto the alum adjuvant (Fig. 1B). We detected CpG1018 in both the unbound and bound to alum fractions.

**Fig. 1.**
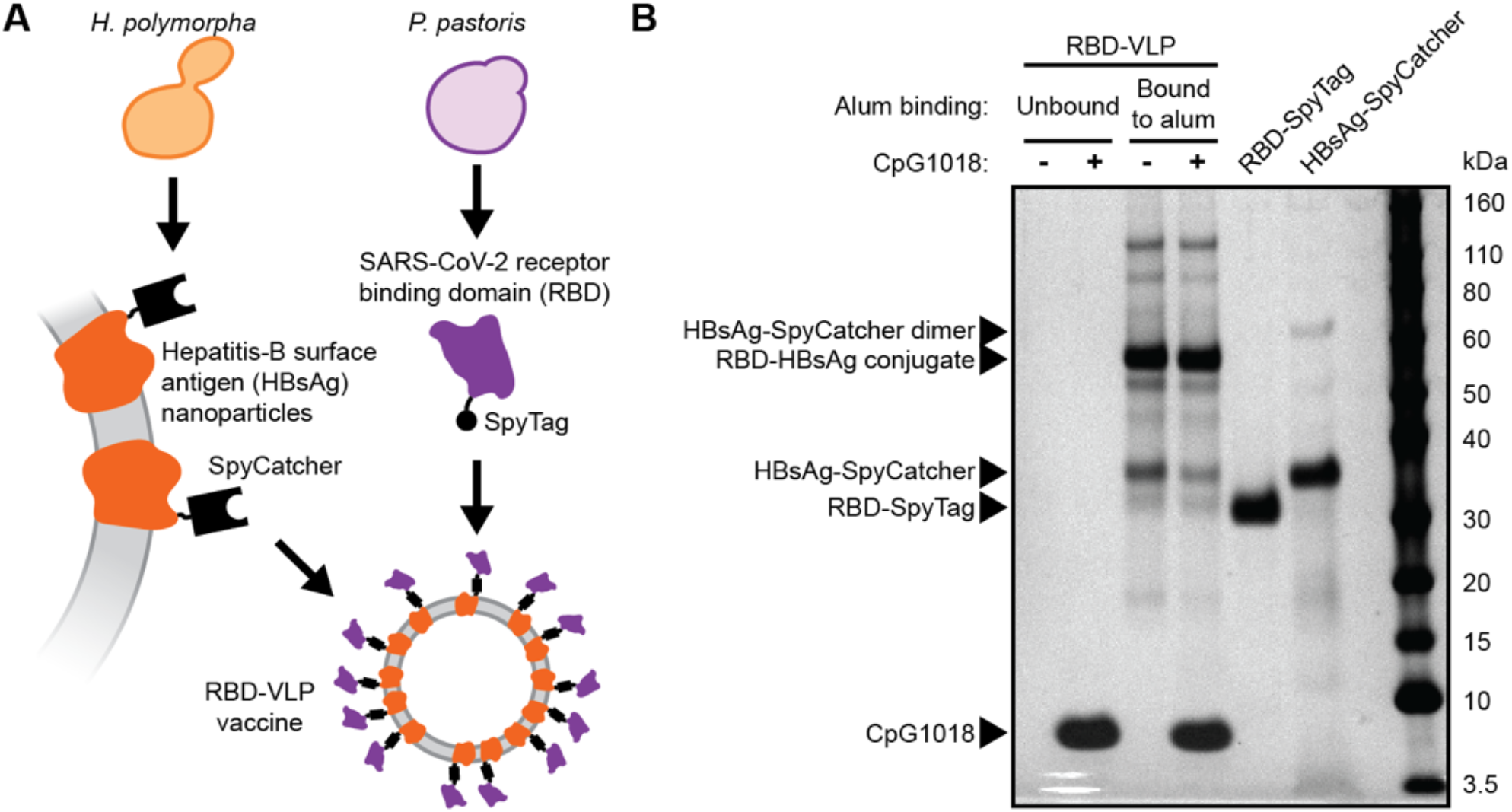
Design and analysis of the RBD-VLP drug product. A) Schematic of protein expression and conjugation. B) Reduced SDS-PAGE analysis of the formulated RBD-VLP vaccine samples. Alum-bound protein antigen (with and without CpG) was separated by centrifugation and desorbed from the alum using an elution buffer combined with heat treatment prior to SDS-PAGE.

We also performed additional analytics on unformulated RBD-VLP antigen to confirm antigenicity and nanoparticle formation. The RBD-VLP antigen exhibited strong binding to the human ACE2 receptor and a known neutralizing antibody CR3022 by biolayer interferometry (Fig. S1A). The large difference in signal observed in this analysis between the RBD-VLP and soluble monomeric RBD-SpyTag confirms the multivalency of the RBD conjugated on the VLP. We also confirmed formation of nanoparticles by electron microscopy (EM) (Fig. S1B-C). These analytics confirmed the structural attributes of the conjugated RBD-VLPs used for non-clinical evaluations here.

### Immunogenicity testing in cynomolgus macaques

To assess the immunogenicity of this RBD-VLP vaccine candidate, we immunized cynomolgus macaques with two doses of either vaccine formulation (alum or alum combined with CpG1018) or a placebo, spaced three weeks apart (Fig. 2A). We assessed spike protein specific antibody titers after each dose (Fig. 2B). We observed full seroconversion and high antibody titers for both RBD-VLP vaccine formulations. Interestingly, formulation with only alum elicited significantly higher antibody titers. We assessed the neutralizing activity of the sera against a SARS-CoV-2 pseudovirus and observed high titers of neutralizing antibodies (Fig. 2C). Formulation with only alum appeared to yield higher neutralization, but was not significantly higher (p>0.1, Kolmogorov-Smirnov test) than the vaccine formulated with both alum and CpG1018. Neutralization of SARS-CoV-2 pseudovirus correlated well with spike protein specific antibody titer (Fig. S2A).

**Fig. 2.**
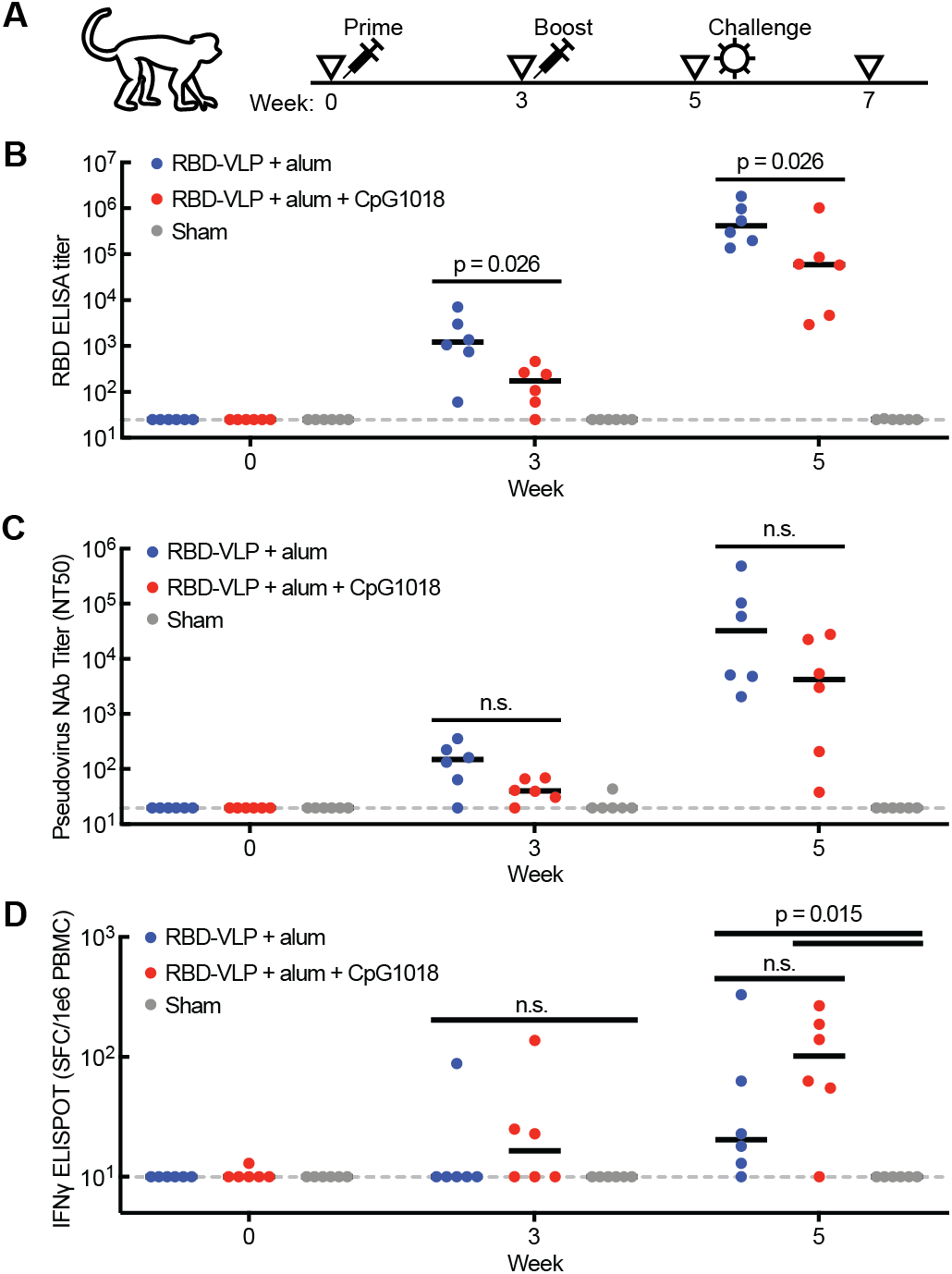
Humoral and cellular immune response to the RBD-VLP vaccine. A) Design of the non-human primate study in cynomolgus macaques. B) Titers of RBD-specific antibody binding in animal sera. C) Titers of SARS-CoV-2 pseudovirus neutralizing antibody in animal sera. D) Expression of IFNγ from cells stimulated with S1 protein peptides. Statistical significance was determined by a Kolmogorov-Smirnov test. N.S. = not significant (p > 0.1). Black bars represent median values. Dotted gray line represents limit of detection.

We also assessed the neutralizing activity of sera from week 5 against SARS-CoV-2 pseudoviruses with spike proteins from several circulating variants of concern (Fig. S2B). Sera from animals immunized with RBD-VLP and alum exhibited similar neutralizing activities against D614G and B.1.1.7 variants, but significantly less neutralization against B.1.351 (~25x, geometric mean).

To determine why the formulation with only alum elicited higher antibody titers, we assessed the antigenicity for retained samples of the RBD-VLP in both vaccine formulations used. We observed ~30% less binding to human ACE2 for the alum and CpG1018 formulation compared to the formulation with alum only (competitive ELISA) (Fig. S2C). The formulation with alum and CpG1018 at the concentrations used here appears to have altered the antigenicity of the RBD-VLP antigen, potentially leading to a reduced humoral immune response.

After assessing the humoral response to the RBD-VLP vaccine candidate, we profiled the cellular immune response. We assessed expression of IFNγ in cells stimulated with peptides from the S1 region of the spike protein, which includes the RBD (Fig. 2D). We observed significant IFNγ expression in cells from both the alum formulation and the alum and CpG1018 formulation after two doses. The cellular response appeared stronger with the alum and CpG1018 co-formulation, consistent with previous reports on the influence of CpG1018 as an adjuvant in vaccines (*29*), but the effect was not significant (p>0.4, Kolmogorov-Smirnov test).

### Challenge with SARS-CoV-2

To assess if the RBD-VLP vaccine could protect animals from infection, we challenged all animals with SARS-CoV-2 two weeks after the second immunization (Fig. 2A). We monitored the course of infection for two weeks by measuring titers of subgenomic RNA (sgRNA) in nasal swabs and bronchoalveolar lavage supernatants. Both formulations of the RBD-VLP vaccine significantly reduced the levels of detected sgRNA in the upper respiratory tract (Fig. 3A), and no sgRNA was detected after day 4 post challenge from the group immunized with RBD-VLP formulated with alum (Fig. 3B). Both RBD-VLP formulations exhibited nearly complete protection from viral infection in the lower respiratory tract—sgRNA was detected in bronchoalveolar lavage supernatants from only three animals (Fig. 3C-D). We observed significant correlation between pre-challenge antibody titers and measured sgRNA levels (Fig. 3E-F).

**Fig. 3.**
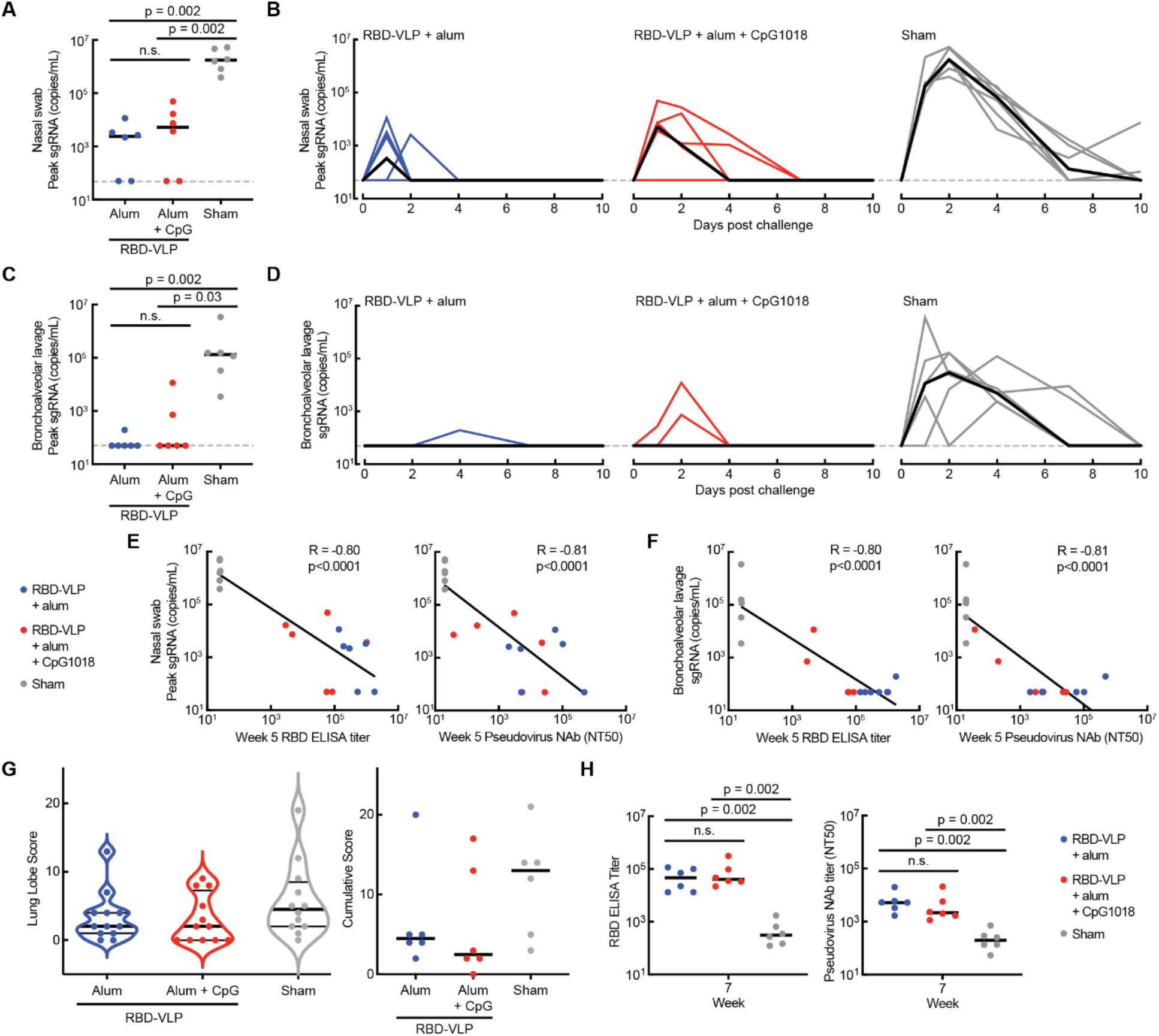
Challenge with SARS-CoV-2. A,C) Peak levels of SARS-CoV-2 sgRNA after challenge for each animal from nasal swabs (A) or bronchoalveolar lavage (C). Statistical significance was determined by a Kolmogorov-Smirnov test. N.S. = not significant (p > 0.1). Black bars represent median values. Dotted gray line represents limit of detection. B,D) Levels of sgRNA after challenge from nasal swabs (B) or bronchoalveolar lavage (D). Each thin line represents one animal. Thick black lines represent median values. E,F) Correlation of RBD-specific antibody titer and pseudovirus neutralizing antibody titer from week 5 animal sera with peak sgRNA levels in nasal swabs (E) and bronchoalveolar lavage (F). R was calculated by Spearman correlation. G) Pathology scores for individual lung samples and cumulative scores for each animal. H) Titers of RBD-specific antibody and SARS-CoV-2 pseudovirus neutralizing antibody in animal sera at week 7 (2 weeks post challenge). Statistical significance was determined by a Kolmogorov-Smirnov test. N.S. = not significant (p > 0.1). Black bars represent median values.

We also assessed lung samples from each animal after challenge with SARS-CoV-2 and scored each sample for histopathologic findings. Animals that received the sham vaccine showed evidence of interstitial inflammation, syncytial cells, and type II pneumocyte hyperplasia in the lung, consistent with SARS CoV-2 replication and a higher median cumulative pathology score (13) as compared to immunized animals (median 2.5-4.5) (Fig. 3G).

Finally, we measured RBD-specific and pseudovirus neutralizing antibody titers after challenge. Animals that received the RBD-VLP vaccine candidate had higher antibody levels than animals that were not immunized pre-challenge (Fig. 3H). Notably, post-challenge levels of humoral immunity conferred by the vaccine were not significantly different between the two vaccine formulations (p>0.4, Kolmogorov Smirnov test). These results together suggest that the RBD-VLP vaccine candidate can protect from SARS-CoV-2 infection.

## Discussion

In this study, we report the design of an RBD-VLP based protein subunit vaccine candidate for COVID-19, currently manufactured in existing production facilities at the Serum Institute of India. Both protein components are produced in yeast, making this vaccine a promising, low-cost intervention for LMICs.

We tested the vaccine in cynomolgus macaques, which have been shown to be equivalent to rhesus macaques as a model for SARS-CoV-2 infection (*30*). The RBD-VLP vaccine elicited high titers of neutralizing antibodies (>10^4^) that were above the range of protection for other licensed vaccines in non-human primates (*31, 32*). The vaccine also conferred protection from viral loads in the upper and lower respiratory tract upon challenge with SARS-CoV-2.

CpG adjuvants with HBsAg vaccines have been shown to boost the Th1 cell response to complement the Th2 cell response induced with alum alone (*29, 33, 34*). In agreement with these prior reports, we did observe a higher Th1 cell response to the RBD-VLP formulated with both alum and CpG1018 compared to alum only. Interestingly, we observed a reduced humoral immune response when the vaccine was formulated with alum and CpG1018. We attributed this difference to a reduction in antigenicity of the RBD; inclusion of CpG may impact the structural integrity of the RBD, leading to altered antigenicity. We have previously observed similar inconsistent immunogenicity of monomeric SARS-CoV-2 RBD when formulated with a CpG adjuvant (*24*). Another multimeric RBD vaccine has reported robust humoral responses in nonhuman primates with the same dose of CpG1018 (*35*), but in that instance, the vaccine was formulated 30 minutes prior to injection. This discrepancy suggests that a high dose of CpG1018 may impact the antigenicity of the RBD when stored for weeks or months at 4°C prior to use. Reduced dosing of CpG1018 could lessen the impact on the stability of the RBD antigen while still eliciting balanced Th1 and Th2 cellular responses (*36*). This question warrants further study to optimize the concentrations of this co-adjuvant in formulations of vaccine candidates to balance the stability of the RBD-VLP antigen with the beneficial effects of the alum and CpG1018 adjuvants.

Another potential challenge posited for CoVID-19 and interventions directed towards SARS-CoV-2 is the possibility of antibody-dependent enhancement (ADE) or vaccine-associated enhanced disease (*37*). Such effects could present safety concerns for novel vaccines (*38*). Both vaccine candidates tested here with alum or CpG and alum as adjuvants did not show evidence for disease enhancement following vaccination and subsequent challenge. This outcome is consistent with evidence that passive immunization with human antibodies directed against SARS-CoV-2 have not exhibited disease enhancement in mice and monkeys (*39*), and provides support that formulations of the vaccine here with alum alone could provide protective immunity without disease enhancement.

The report of this vaccine comes at a befitting time in the COVID-19 pandemic. Variants of SARS-CoV-2 with increased transmissibility continue to drive outbreaks of COVID-19 in LMICs. Vaccines based on mRNA have been shown to be effective against emerging variants (*40, 41*), but remain largely inaccessible in southeast Asia and Africa due to high costs and cold chain requirements. Responses induced by currently approved protein-based vaccines based on the Wuhan-Hu-1 strain, on the other hand, have so far been less effective in neutralizing activity exhibited with sera against circulating variants (*42*). Encouragingly, vaccines based on the RBD alone effectively boost an immune response originally generated against a full-length spike protein trimer (*43*). There is growing interest, therefore, in the utility of RBD-based vaccine boosters to provide immunity against emerging variants of SARS-CoV-2 that may escape antibody responses induced by vaccination against the Wuhan-Hu-1 or D614G spike protein.

The RBD-VLP vaccine candidate presented here is modular, and could be manufactured and formulated with RBD antigens from emerging variants or with engineered immunogenicity (*24*). We reported previously that the RBDs from the B.1.1.7 and B.1.351 variants can be manufactured from the same microbial strains and fermentation processes (*28*). A mosaic vaccine against multiple strains of SARS-CoV-2 could be simply produced by conjugation of multiple RBD antigens to the HBsAg-VLP. This approach could also be tailored to generate multivalent designs for a panel of SARS-CoV-2 variants or for broadly targeting betacoronaviruses (*22*). This modular design of protein antigens paired with existing large-scale manufacturing capacity in yeast demonstrates a vaccine platform that could keep pace with an evolving COVID-19 pandemic, and provide better access to COVID-19 vaccines around the globe.

## Materials and Methods

### Production of vaccine samples

Hepatitis B Surface Antigen (HBsAg) SpyCatcher nanoparticles and RBD-SpyTag protein were produced by yeast fermentation in a GMP process at the Serum Institute of India Pvt. Ltd. Samples were stored at 4°C for several weeks before immunization.

### Sodium dodecyl sulfate polyacrylamide gel electrophoresis (SDS-PAGE)

Formulated RBD-VLP samples were centrifuged at 4000 g for 10 min at 4°C. The supernatant with unbound protein was transferred to another tube without disturbing the pellet and prepared for SDS-PAGE in 50 mM dithiothreitol (Thermo scientific, A39255), 1X LDS sample buffer (Invitrogen, NP0007), and 20 mM iodoacetamide (Thermo scientific, A39271), and heated at 95°C for 15 min. The alum-bound protein in the pellet was desorbed by resuspending the pellet in an elution buffer (0.4 M sodium phosphate, 50 mM dithiothreitol, 1X LDS sample buffer, and 20 mM iodoacetamide) and heating the samples at 95°C for 15 min. The desorbed samples were then centrifuged at 4000 g for 10 min at 4°C to obtain desorbed protein in the supernatant for SDS-PAGE. RBD-SpyTag and HBsAg-SpyCatcher controls were prepared for SDS-PAGE similar to unbound protein sample. Approximately 0.1 μg of each sample was loaded on 4-12% Bis-Tris gel (Invitrogen, NP3022) run for 50 min at 150 V in 1X MES-SDS running buffer (Invitrogen, NP0002). Novex™ Sharp Pre-stained protein standard (Invitrogen, LC5800) was used as marker to estimate molecular weight of the proteins. The gel was stained using Pierce Silver Stain kit (Thermo, 24612) as per manufacturer’s protocol and imaged using ProteinSimple FluorChem E imaging system.

### Competitive ELISA by ACE2-Fc binding

Stock adjuvanted RBD-VLP samples were first incubated with a blocking buffer, then 2-fold serial dilutions were made followed by incubation with 0.02 mcg/mL ACE2-Fc receptor overnight on a plate rotator at room temperature. Corresponding blank samples were prepared using blocking buffer alone, while the saturated samples contained ACE2-Fc receptor. Samples were centrifuged the next day and the supernatant containing unbound ACE2-Fc was transferred to a 96-well plate coated with 1 mcg/mL of RBD. The plate was incubated for two hours at 25°C, washed, and the amount of bound ACE2-Fc on the plate was detected with a horseradish peroxidase conjugated secondary antibody using a tetramethylbenzidine substrate. The % relative ACE2-Fc binding of samples was determined from the OD_450_ values using the parameters obtained from a 4-point logistic fit of the standard run on each plate.

### Production of unformulated drug substance

Hepatitis B Surface Antigen (HBsAg) SpyCatcher nanoparticles were produced in a GMP process at the Serum Institute of India. RBD-SpyTag protein was expressed in shake flask culture of *Pichia pastoris* and purified as described previously (*24, 28*). HBsAg-SpyCatcher and RBD-SpyTag were conjugated by incubation overnight at 4°C in 20 mM sodium phosphate, 150 mM NaCl, pH 8 buffer, with a 1:1.5 HBsAg:RBD molar ratio. Excess RBD was removed with a 100 kDa molecular weight cutoff Amicon^®^ Ultra-4 centrifugal filter (Millipore).

### Biolayer interferometry

Biolayer interferometry was performed using the Octet Red96 with Protein A (ProA) biosensors (Sartorius ForteBio, Fremont, CA), which were hydrated for 15 min in kinetics buffer prior to each run. Kinetics buffer comprising 1X PBS pH 7.2, 0.5% BSA, and 0.05% Tween 20 was used for all dilutions, baseline, and disassociation steps. CR3022 and ACE2-Fc were used in the assay at concentrations of 2 and 10 μg/mL, respectively. Samples were loaded in a 96-well black microplate (Greiner Bio-One, Monroe, NC) at starting concentrations of 15 and 10 μg/mL, respectively. Seven 1:1 serial dilutions and a reference well of kinetics buffer were analyzed for each sample. Association and dissociation were measured at 1000 rpm for 300 and 600 sec, respectively. Binding affinity was calculated using the Octet Data Analysis software v10.0 (Pall ForteBio), using reference subtraction, baseline alignment, inter-step correction, Savitzky-Golay filtering, and a global 1:1 binding model.

### Negative-stain electron microscopy

Solution with conjugated RBD-VLP nanoparticles (7 μL) was incubated on a 200 meshes copper grid coated with a continuous carbon film for 60 seconds. Excess liquid was removed, and the film was incubated in 10 μL of 2% uranyl acetate. The grid was dried at room temperature and mounted on a JEOL single tilt holder in the TEM column. The specimen was cooled by liquid nitrogen, and imaged on a JEOL 2100 FEG microscope with a minimum dose to avoid sample damage. The microscope was operated at 200 kV with magnification at 10,000-60,000x. Images were recorded on a Gatan 2Kx2K UltraScan CCD camera.

### Immunization of non-human primates

18 cynomolgus macaques were randomly allocated to groups. Animals were housed at Bioqual, Inc. (Rockville, MD). On day 0, groups of animals were immunized with: 1) RBD-HBSAg VLP containing 5μg total protein adjuvanted with 500 μg Alum, 2) RBD-HBSAg VLP containing 5 μg protein adjuvanted with 500 μg of Alum + 1500 μg CpG, or 3) sham. Animals were administered an identical boost immunization of the regimens as indicated above at week 3. At week 6, all animals were challenged with 1 x 10^5^ TCID50 SARS-CoV-2 via the intranasal and intratracheal routes (2 mL total volume per animal; 1 mL intranasal + 1 mL intratracheal). The challenge stock was derived from a single passage of a SARS-CoV-2 2019 USA-WA1/2020 isolate (BEI Resources, NR-53780).

### ELISA assays

Binding antibodies were quantified by ELISA essentially as previously described (*44*). Briefly, 96-well plates were coated with 1μg/ml SARS-CoV-2 S protein (Sino Biological) in 1X DPBS and incubated overnight at 4°C. Plates were then washed once with wash buffer (0.05% Tween 20 in 1 X DPBS) and blocked for 2-3 h with 350 μL Casein block/well at room temperature. After block, serial dilutions of serum diluted in casein block were added to wells and plates were incubated for 1 h at room temperature. Plates were then washed three times, a 1μg/ml dilution of anti-macaque IgG HRP (Nonhuman Primate Reagent Resource) was added to wells, and plates were incubated at room temperature in the dark for 1 h. Plates were again washed three times, then 100 μL of SeraCare KPL TMB SureBlue Start solution was added to each well. Plate development was halted with the addition of 100 μL SeraCare KPL TMB Stop solution per well. A VersaMax microplate reader was used to record the absorbance at 450nm. For each sample, endpoint titer was calculated in Graphpad Prism software, using a four-parameter logistic curve fit to calculate the reciprocal serum dilution that yields an absorbance value of 0.2 at 450nm. Log_10_ endpoint titers are reported.

### Pseudovirus neutralization assays

SARS-CoV-2 pseudoviruses were generated essentially as described previously (*44, 45*). Briefly, HEK293T cells were co-transfected with: 1) the packaging plasmid psPAX2 (AIDS Resource and Reagent Program), 2) luciferase reporter plasmid pLenti-CMV Puro-Luc (Addgene), and 3) an S protein-expressing plasmid, pcDNA3.1-SARS CoV-2 SΔCT using lipofectamine 2000 (ThermoFisher). This approach was used to generate pseudoviruses specific to SARS-CoV-2 variant strains, including WA1/2020 strain (Wuhan/WIV04/2019, GISAID accession ID: EPI_ISL_402124), D614G mutation, B.1.1.7 variant (GISAID accession ID: EPI_ISL_601443), and B.1.351 variant (GISAID accession ID: EPI_ISL_712096). To purify pseudoviruses, supernatants were collected 48 h post-transfection, centrifuged, then passed through a 0.45 μm filter. To determine the neutralization activity of the serum samples from nonhuman primates, HEK293T-hACE2 cells were seeded in 96-well tissue culture plates at a density of 1.75 x 10^4^ cells/well and incubated overnight. Serial dilutions of heat inactivated serum samples were prepared and mixed with 50 μL of indicated pseudovirus. This mixture was incubated at 37°C for 1 h before adding to seeded HEK293T-hACE2 cells. 48 h after infection, cells were lysed in Steady-Glo Luciferase Assay Reagent (Promega) according to the manufacturer’s instructions.

SARS-CoV-2 neutralization titers were defined as the sample dilution at which a 50% reduction in relative light unit (RLU) was observed relative to the average of the virus control wells.

### ELISPOT assays

ELISPOT assays were performed essentially as described previously (*44*). Plates were coated with mouse anti-human IFN-γ antibody (BD Pharmigen) at 5 μg/well and incubated at 4°C overnight. After incubation, plates were washed with wash buffer (1 x DPBS with 0.25% Tween20), then blocked with R10 media (RPMI with FBS and penicillin-streptomycin) for 1-4 h at 37°C. SARS-CoV-2 S1 peptides were prepared and plated at 1 μg/well. 200,000 cells/well were added to the plate and incubated for 18-24 h at 37°C. Plates were then washed with wash buffer (MilliQ water with DPBS & Tween20) and incubated for 2 h at room temperature with polyclonal rabbit anti-human IFN-γ biotin from U-Cytech at 1 μg/mL. The plates were washed again, then incubated for 2 h at room temperature with Streptavidin-alkaline phosphatase from Southern Biotech at 2 μg/mL. A final wash step was followed by the addition of Nitro-blue Tetrazolium Chloride/5-bromo-4-chloro 3 ‘indolyphosphate p-toludine salt (NBT/BCIP chromogen) substrate solution for 7 min. The chromogen was discarded and the plates were washed with water and dried for 24 h protected from light. Plates were scanned and counted on a Cellular Technologies Limited Immunospot Analyzer.

### Viral load assays

SARS-CoV-2 E gene subgenomic RNA (sgRNA) was assessed by RT-PCR using primers and probes essentially as previously described (*44*). To generate a standard, a fragment of the subgenomic E gene was synthesized and cloned into a pcDNA3.1+ expression plasmid using restriction site cloning (Integrated DNA Techonologies). The insert was *in vitro* transcribed to RNA using the AmpliCap-Max T7 High Yield Message Maker Kit (CellScript). Log-fold dilutions were prepared for RT-PCR assays, ranging from 1×10^10^ copies to 1×10^-1^ copies. To quantify viral loads in respiratory tract tissues, samples of bronchoalveolar lavage (BAL) fluid and nasal swabs (NS) were analyzed. RNA extraction was performed on a QIAcube HT using the IndiSpin QIAcube HT Pathogen Kit according to manufacturer’s specifications (Qiagen). Standards described above and extracted RNA from nonhuman primate samples were reverse transcribed using SuperScript VILO Master Mix (Invitrogen), using the cycling conditions specified by the manufacturer, 25° C for 10 min, 42° C for 1 h, and then 85° C for 5 min. A Taqman custom gene expression assay (Thermo Fisher Scientific) was designed using the sequences targeting the E gene sgRNA. The sequences for the custom assay were as follows: forward primer, sgLeadCoV2.Fwd: CGATCTCTTGTAGATCTGTTCTC, E_Sarbeco_R: ATATTGCAGCAGTACGCACACA, E_Sarbeco_P1 (probe): VIC-ACACTAGCCATCCTTACTGCGCTTCG-MGB. Reactions were carried out in duplicate for samples and standards on the QuantStudio 6 and 7 Flex Real-Time PCR Systems (Applied Biosystems) with the following thermal cycling conditions: initial denaturation at 95°C for 20 s, followed be 45 cycles of 95° C for 1 s and 60° C for 20 s. Standard curves were used to calculate sgRNA copies per mL of BAL fluid or per swab; the quantitative assay sensitivity was 50 copies per mL or per swab.

### Histopathology

At time of fixation, lungs were suffused with 10% formalin to expand the alveoli. All tissues were fixed in 10% formalin and blocks sectioned at 5 μm. Slides were baked for 30-60 min at 65 degrees, deparaffinized in xylene, rehydrated through a series of graded ethanol to distilled water, then stained with hematoxylin and eosin (H&E). Blinded histopathological evaluation was performed by a board-certified veterinary pathologist (AJM). Two lung lobes (1 section from the right and left caudal lung lobes) were assessed and scored (1-4) for each of the following lesions: 1) Interstitial inflammation and septal thickening 2) Eosinophilic interstitial infiltrate 3) Neutrophilic interstitial infiltrate 4) Hyaline membranes 5) Interstitial fibrosis 6) Alveolar infiltrate, macrophage 7) Alveolar/Bronchoalveolar infiltrate, neutrophils 8) Syncytial cells 9) Type II pneumocyte hyperplasia 10) Broncholar infiltrate, macrophage 11) Broncholar infiltrate, neutrophils 12) BALT hyperplasia 13) Bronchiolar/peribronchiolar inflammation 14) Perivascular, mononuclear infiltrates and 15) Vessels, endothelialitis. Each feature assessed was assigned a score of 0= no significant findings; 1=minimal; 2= mild; 3=moderate; 4=marked/ severe.

## Acknowledgements

The authors thank the Koch Institute’s Robert A. Swanson (1969) Biotechnology Center for technical support. We thank Jake Yalley-Ogunro, Jeanne Muench, Alex Granados, and John Harrison for assistance with the immunization phase of the non-human primate study. The following reagent was deposited by the Centers for Disease Control and Prevention and obtained through BEI Resources, NIAID, NIH: SARS-Related Coronavirus 2, Isolate USA-WA1/2020, NR-53780.This work was funded by the Bill and Melinda Gates Foundation (Investment IDs INV-002740, INV-006131, and INV-027406). This study was also supported in part by the Koch Institute Support (core) Grant P30-CA14051 from the National Cancer Institute. L.H.T. is an NIH T32 Postdoctoral Fellow supported by the Multidisciplinary AIDS Training Program (Grant # T32 AI007387). The content is solely the responsibility of the authors and does not necessarily represent the official views of the NCI, the NIH, Gates MRI, or the Bill and Melinda Gates Foundation.

## Author contributions

N.C.D., L.H.T., S.R.A., J.C.L., and D.H.B. conceived and planned experiments. K.K., S.B., and O.S.K. analyzed vaccine samples. N.C.D., S.R.A., C.A.N., and L.B. analyzed unformulated drug substance. D.Y. performed electron microscopy. L.H.T. coordinated and analyzed animal experiments. A.C., K.M., N.B.M., J.Y., A.C., V.M.G., F.N., and S.P. performed immunologic and virologic assays. A.J.M. performed the histopathology. Z.F., L.P., R.B., J.V., E.T., A.C., H.A., and M.G.L. led clinical care of non-human primates. D.C. and J.M.S. coordinated and managed resource allocations and planning for experimental studies. N.C.D., L.H.T., S.R.A., J.C.L., and D.H.B. wrote the manuscript. D.H.B., J.C.L., S.B., D.B.V., S.B.J., and H.K. designed the experimental strategy and reviewed analyses of data. All authors reviewed the manuscript.

---

**Fig. S1.**
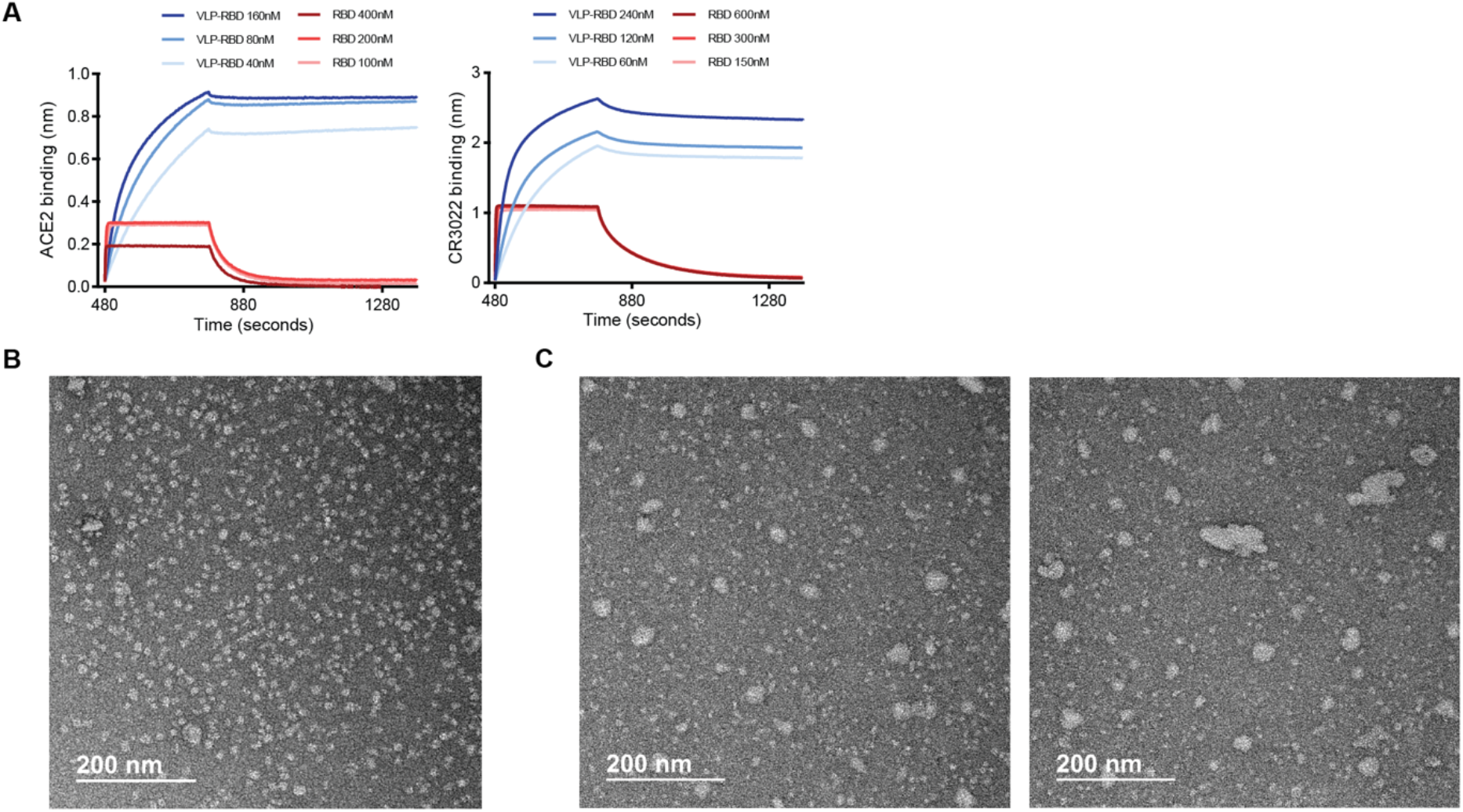
Analysis of representative samples of RBD-VLP antigen. A) Biolayer interferometry of binding to human ACE2-Fc protein, and CR3022 neutralizing antibody. RBD-VLP conjugated protein is shown in blue. RBD-spytag monomer is shown in red. B-C) Negative stain electron microscopy of unconjugated HBsAg-VLP (B) and conjugated RBD-VLP (C).

**Fig. S2.**
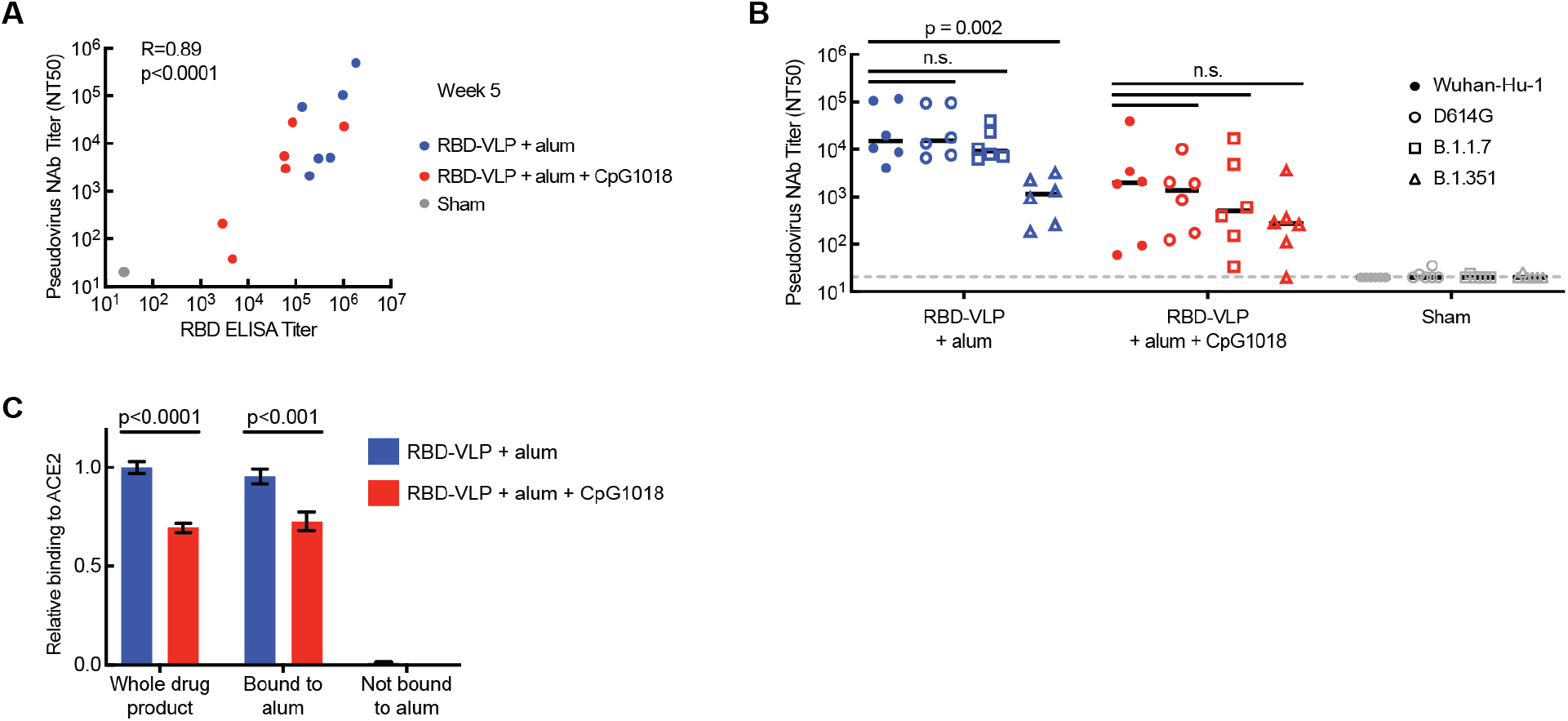
Further analysis of the humoral immune response. A) Correlation of RBD-specific antibody titer and pseudovirus neutralizing antibody titer from week 5 animal sera. R was calculated by Spearman correlation. B) Titers of neutralizing antibodies to SARS-CoV-2 variant pseudoviruses in animal sera. Statistical significance was determined by a Kolmogorov-Smirnov test. N.S. = not significant (p > 0.1). Black bars represent median values. Dotted gray line represents limit of detection. C) Relative binding of adjuvanted RBD-VLP to ACE2-Fc by competitive ELISA. Alum-bound protein was separated to bound and unbound only fractions by centrifugation. Error represents standard deviation after two independent measurements.

